# Beyond phylogeny: Genome-wide DNA sequence patterns suggest DNA physical properties associated with thermal adaptation in extremophile microbes

**DOI:** 10.64898/2026.06.16.732181

**Authors:** Monireh Safari, Lila Kari

## Abstract

Temperature is a fundamental constraint on biological systems, yet how it is reflected in genome sequence organization remains unclear. Here, we show that genome-wide distributions of short DNA sequences contain a robust signal of thermal adaptation that is largely independent of phylogeny. Using Structural Topic Modelling (STM), a machine-learning approach for identifying groups of co-occurring sequence motifs, we analyze canonical 6-mer and 9-mer frequency profiles of bacterial and archaeal genome proxies (randomly sampled genomic regions) and identify motif families systematically associated with thermophiles and psychrophiles. In bacterial thermophiles, the identified motif families are dominated by highly specific, overrepresented and co-occurring C- and G-stacked hexamers, and a distinct family of CG-periodic hexamers recurring across multiple temperature comparisons. In contrast, bacterial psychrophile-associated motifs are dominated by low-complexity A-, T-, and AT-run hexamers. Thermophilic archaea generally exhibit a distinct CTAG-centred hexamer family, suggesting that different domains may adapt to similar environmental constraints through different sequence-level solutions. However, this domain-level contrast is not absolute: in a targeted analysis of two thermophilic bacterium–archaeon pairs, we find unusually similar frequencies of all the STM-identified thermophile-associated hexamer families, suggesting that shared high-temperature environments can, in specific cases, partially override phylogenetic divergence. Notably, the identified motif families constitute only a small and highly selective subset of the vast space of possible G+C-rich or A+T-rich sequences. This indicates that thermal adaptation is associated with specific sequence architectures rather than broad shifts in nucleotide composition. Accordingly, the observed signal cannot be explained by overall base composition alone, but instead arises from structured combinations and positional arrangements of nucleotides within short sequence contexts. Related motif families are recovered at both *k* = 6 and *k* = 9, indicating that the signal reflects systematic shifts in genome-wide sequence organization rather than isolated sequence motifs. These patterns are consistent with known sequence-dependent DNA physical properties documented in biochemical and biophysical studies, including differences in base-stacking interactions and conformational flexibility. Together, our results suggest that genome-wide sequence organization reflects sequence-dependent DNA physical properties associated with thermal adaptation, revealing a previously underappreciated physical layer of genomic information beyond phylogenetic history.

## 1 Introduction

Genome-wide DNA sequence composition encodes a robust and previously underexplored signal of thermal adaptation that sometimes transcends phylogenetic boundaries. Here, we show that higher-order short DNA motifs (*k*-mers, for *k* = 6 and *k* = 9), considered collectively across the genome, systematically distinguish thermophilic and psychrophilic microorganisms and reveal patterns consistent with sequence-dependent DNA physical properties. These findings suggest that thermal adaptation is associated with pervasive higher-order sequence constraints acting across the genome rather than with isolated genes or simple shifts in nucleotide composition alone.

Microorganisms thriving at extreme temperatures, thermophiles and hyperthermophiles with optimal growth above 45◦C and 80◦C, and psychrophiles with optima below 20◦C, provide a natural setting to study how environmental constraints shape genomic organization [3, 7]. Temperature affects the stability and flexibility of biomolecules, and signatures of thermal adaptation have been identified at multiple levels, including proteins, RNA structures, and nucleotide composition [30, 9]. However, the genomic basis of thermal adaptation remains incompletely understood.

A central question is whether genomic base composition reflects adaptation to temperature. Although G+C content confers greater thermodynamic stability, comparative analyses have shown that there is no consistent, genome-wide correlation between G+C content and optimal growth temperature, once phylogeny is taken into account [6, 14, 28]. This being said, strong correlations have been observed in structural RNAs, where increased G+C content stabilizes functional secondary structures at elevated temperatures [6]. Together, these observations indicate that temperature can affect nucleic acid sequence, but that genome-wide nucleotide composition alone (e.g. G+C content) does not fully explain how this environmental component is encoded in DNA.

In this context, DNA biophysics provides a framework for understanding why thermal adaptation may be reflected in local sequence context rather than in base composition alone. For example, it was observed that duplex stability is governed primarily by base-stacking interactions rather than base composition alone [21, 22, 29], and also that DNA mechanical properties depend on sequence context extending to tetra- and hexanucleotide scales [16, 11]. Genome-scale studies further show that these sequence-dependent physical properties are non-randomly distributed [2, 1]. Together, these findings suggest that thermal adaptation may be reflected in higher-order sequence organization rather than nucleotide composition alone.

Alignment-free approaches based on *k*-mer frequencies capture such higher-order context at the genome scale. Lower-order patterns (e.g., *k* = 3), which largely reflect codon usage and nucleotide composition, have been studied extensively [13, 20]. In particular, it was shown that *k*-mer features can predict optimal growth temperature and detect environmental signals in extremophile genomes [23, 13, 20]. However, these approaches detected environment-associated signals at the genome level, and did not identify specific DNA motifs driving these similarities. Moreover, these studies did not offer interpretations of the similarity of *k*-mer frequency profiles of same environment-type microbial genomes.

Here, we address this gap by proposing a novel application of Structural Topic Modelling (a machine learning method) to genome proxy sequences, in order to identify specific DNA motifs driving their similarities. STM is a natural language processing (NLP) method used to automatically discover hidden themes (topics) across large collections of unstructured text. Focusing on hexamers (*k* = 6) and longer motifs (*k* = 9), we use STM to identify latent motif classes whose prevalence is associated with thermal adaptation while controlling for phylogenetic effects. In bacteria, thermophile-associated motifs are enriched in C- or G-stacked motifs and CG-periodic motifs, whereas psychrophile-associated motifs are dominated by A-, T-, or AT-run motifs. Importantly, this signal is not reducible to overall G+C content. Indeed, genomes with similar base composition can exhibit markedly different motif distributions. Instead, discrimination arises from structured combinations and positional arrangements of nucleotides within short sequence contexts, consistently recovered across *k* and across taxa. In thermophilic archaea, we identify a distinct CTAG-centred motif family, suggesting that different domains may adapt to similar environmental constraints through different sequence-level solutions.

The main contributions of this paper are:

1. A novel application of Structural Topic Modelling to microbial *k*-mer frequency profiles, revealing genome-wide motif families associated with thermal adaptation in bacteria and archaea.
2. A robust and reproducible computational pipeline, including multi-run consensus stabilization and genus-level phylogenetic control, to ensure that identified motif families are not artifacts of randomness in model initialization.
3. A connection between thermophile- and psychrophile-associated motif families, and sequence classes with known DNA structural and mechanical properties, suggesting a physical interpretation of genome-wide thermal adaptation signals.

Taken together, our results show that genome-wide distributions of higher-order DNA motifs encode a strong, phylogeny-independent signature of thermal adaptation consistent with established principles of DNA thermodynamics and mechanics. This provides a physically grounded interpretation of *k*-mer-based genomic signatures.

## 2 Methods

This section details the dataset used in this study and the method employed to identify extreme environment-type adaptation signals in genomic signatures.

### 2.1 Datasets

In this study, we used the *Temperature Dataset* introduced in [13, 20], composed of 598 genomes, including 369 bacteria and 229 archaea genomes. Three temperature categories are defined based on the optimal temperature for growth (OTG): Psychrophiles (OTG of *<*20°C), Mesophiles (OTG of 20–45°C), and Thermophiles (OTG of *>*45°C) based on the categories introduced in [12]. The taxonomic diversity of the dataset across both domains and all three temperature categories is summarized in Table 1. An overview of the taxonomic composition across domains and phyla is shown in Figure 1, and per-domain breakdowns at the phylum–class level are provided in the Supplementary Materials, Section A.

**Table 1:**
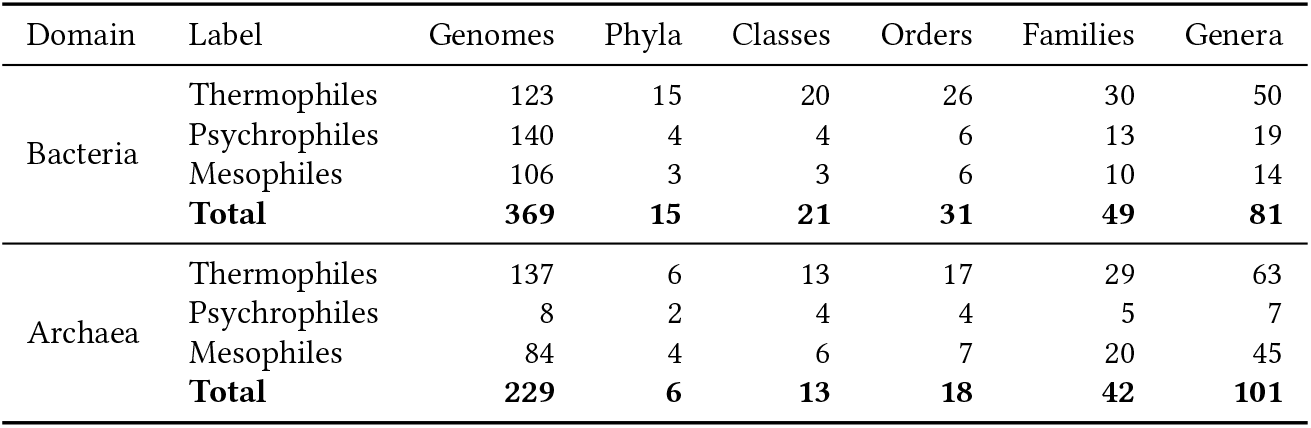
Taxonomic diversity of the Temperature Dataset, including polyextremophilic genomes, across bacteria and archaea.

| Domain | Label | Genomes | Phyla | Classes | Orders | Families | Genera |
| --- | --- | --- | --- | --- | --- | --- | --- |
| Bacteria | Thermophiles | 123 | 15 | 20 | 26 | 30 | 50 |
|  | Psychrophiles | 140 | 4 | 4 | 6 | 13 | 19 |
|  | Mesophiles | 106 | 3 | 3 | 6 | 10 | 14 |
|  | **Total** | **369** | **15** | **21** | **31** | **49** | **81** |
| Archaea | Thermophiles | 137 | 6 | 13 | 17 | 29 | 63 |
|  | Psychrophiles | 8 | 2 | 4 | 4 | 5 | 7 |
|  | Mesophiles | 84 | 4 | 6 | 7 | 20 | 45 |
|  | **Total** | **229** | **6** | **13** | **18** | **42** | **101** |

**Figure 1:**
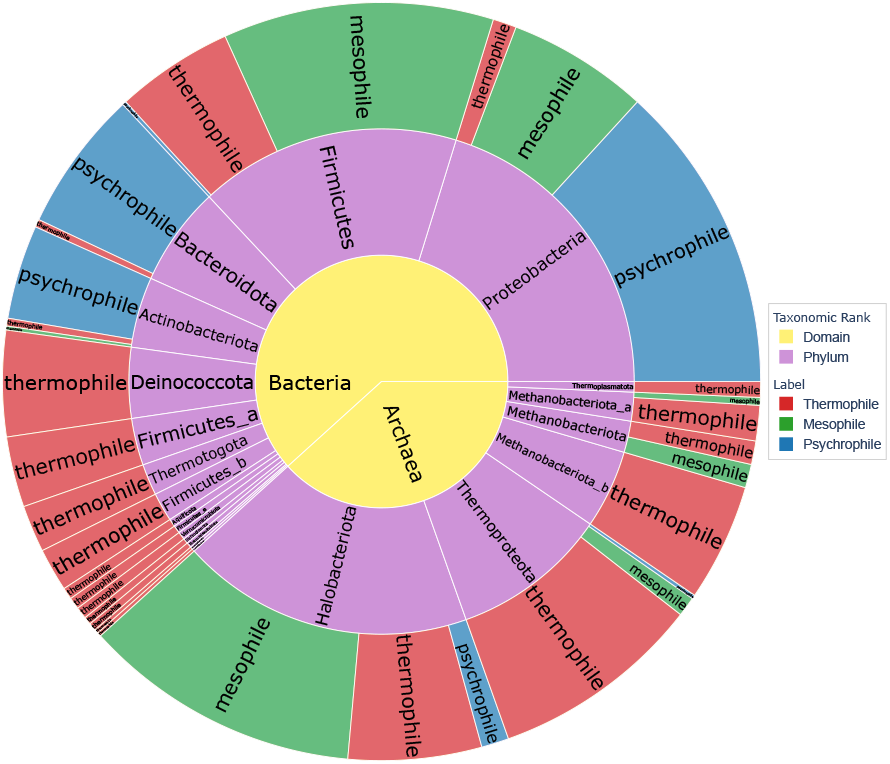
Taxonomic diversity of the Temperature Dataset (598 genomes). The sunburst diagram shows the hierarchical breakdown from domain (inner ring) to phylum (middle ring) to temperature label (outer ring). Arc width is proportional to genome count; colours indicate temperature category: thermophile (red), mesophile (green), and psychrophile (blue).

### 2.2 Genome proxy selection and *k*-mer frequency vector calculation

Following [20], which demonstrated the genome-wide pervasiveness of *k*-mer frequency profiles, we represent each genome by a *genome proxy*. Genome proxy is a composite DNA fragment of fixed length assembled from randomly sampled sub-fragments selected from across the genome. The genomic signature of each genome proxy was calculated as its *k*-mer frequency vector, wherein each entry records the relative frequency of a *canonical k-mer* and its Watson-Crick complement. A canonical *k*-mer is defined as the lexicographically smaller of a *k*-mer and its Watson–Crick complement [20, 13]. For example, if *k* = 6 and a DNA sequence has 1 occurrence of the AACCCC and 3 occurrences of its Watson-Crick complement GGGGTT, then the corresponding vector entry would count 4 occurrences for the calculation of the relative frequency of AACCCC (the canonical hexamer of this pair).

Whereas [20] used a single proxy per genome for classification, our analysis feeds multiple genome proxies into STM, to leverage the fact that having multiple documents per genome increases the corpus size and exposes the model to broader genomic coverage from each genome sequence. We therefore generate *R* = 10 independent proxies per genome by repeating the construction procedure above with 10 different random seeds. Each genome proxy is treated as an independent document in the STM corpus, and its genome of origin is recorded in the document metadata so that genome-level grouping can be recovered at the analysis stage when topic significance is analyzed.

### 2.3 Structural Topic Modelling (STM) of canonical *k*-mer profiles

We utilize Structural Topic Modelling (STM) [19] to discover latent DNA patterns in the canonical *k*-mer profiles of genome proxies, namely patterns that are systematically associated with adaptation to extremely low or extremely high temperatures.

A collection of documents can be described by a mixture of topics. For example, a newspaper article might be 70% politics and 30% economics, while another might be 80% sports and 20% culture. STM is a specialized machine learning method that can infer such topics from the words that tend to co-occur across documents. In our case, STM treats each genome proxy as a document, each distinct canonical *k*-mer as a word, and each “*k*-mer topic” as a collection of specific *k*-mers that co-occur across genome proxies. Note that, throughout this paper, the term “topic” refers to a “*k*-mer topic”, where the value of *k* is understood from the context.

Unlike standard topic models, STM allows the prevalence of each topic to be modelled as a function of genome-level metadata. In this study, we use the temperature category as the main variable of interest and genus as a taxonomic covariate. Thus, STM does not only infer groups of co-occurring *k*-mers; it also estimates whether each group is significantly more prevalent in one temperature category versus the comparison group, after accounting for genus-level taxonomic structure. For example, the model can identify a G+C-rich *k*-mer topic whose prevalence is higher in thermophilic genomes, or an A+T-rich low-complexity topic whose prevalence is higher in psychrophilic genomes.

Moreover, rather than assigning each genome exclusively to one topic, STM assigns every genome proxy a set of weights across all topics, reflecting the degree to which each pattern is present in that genome proxy’s *k*-mer profile. These weights are then used to estimate which topics are associated with the temperature category of the corresponding genome, while accounting for genus. Formally, the weight that a given genome proxy *g* places on each topic is captured in its topic proportion vector 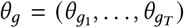, where *θ*_*gt*_ ≥ 0 and 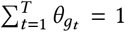; we use *T* for the number of topics instead of the standard notation *K*, to avoid confusion with *k*-mer length. Each genome proxy is then represented as:

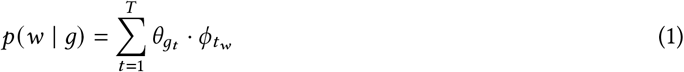

where 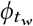 is the probability of *k*-mer *w* under topic *t* .

The prevalence formula includes two covariates. The first is the temperature category label: for bacteria, all three pairwise comparisons are used, namely thermophile–mesophile, psychrophile–mesophile, and thermophile– psychrophile, whereas for archaea only the thermophile–mesophile comparison is included, as the archaeal psychrophile sample is too small for reliable STM fitting (see Table 1). This allows the model to identify topics systematically enriched or depleted in extreme-temperature organisms relative to the comparison group. The second covariate is the genus of each genome, included to account for genus-level taxonomic structure in *k*-mer composition (without this adjustment, inferred topics could reflect taxonomic relatedness rather than temperature-associated sequence patterns). Genera with fewer than two representatives are excluded before model fitting to ensure stable covariate estimation. See Supplementary Materials, Section B for more technical details.

#### 2.3.1 Multi-run consensus and topic stability

Topic models can converge to different local optima depending on the random initialisation, so a single run may not reliably represent the true structure in the data. To obtain robust results, we fit each STM configuration ten times, each time using a different random seed drawn deterministically from a fixed master seed so that the procedure is fully reproducible.

After fitting, we align the topics from runs 2 through 10 to the topics found in run 1. Alignment is performed greedily: for each topic in run 1, we find the topic in the current run whose *k*-mer distribution is most similar to it (measured by cosine similarity), and relabel it accordingly. This ensures that for example “Topic 2” refers to the same latent pattern across all runs before any averaging takes place. Note that the label order of the topics is arbitrary, STM assigns topic indices randomly, so this relabelling does not change the meaning of the inferred topics, only their indices.

Once topics are aligned, we compute two summary quantities. First, we calculate *topic stability* for each topic as the mean pairwise cosine similarity between that topic’s *k*-mer distribution across all pairs of runs. A stability score close to 1 means the topic was recovered consistently in every run; scores below the stability threshold are flagged as unstable. Second, once topics are aligned across all runs, we average both the topic *k*-mer distributions and the proxy topic proportion matrices element-wise across the runs. This means that for each topic, its final *k*-mer profile is the average of what that topic looked like across all runs, and for each genome proxy, its final membership score for each topic is the average across all runs. The result is a single, more stable representation that is less likely to reflect the artifacts of any one random starting point. All subsequent steps use the consensus-averaged 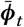 and 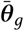, the final *k*-mer profile of each topic and the final proportion vector of each genome proxy (see Equation 1), rather than the output of any individual run.

#### 2.3.2 Identification of driving *k*-mers

For each topic, we want to identify the *k*-mers that are most characteristic of that topic compared to all others, its *driving k-mers*. Following [27], who applied Kullback-Leibler (KL)-divergence-based distinctiveness scoring to nucleotide sequence topic models, we assign each *k*-mer a KL-distinctiveness score. For every *k*-mer in a given topic, the score measures how different that topic’s probability of producing this *k*-mer is from the *most similar* other topic. Using the most similar comparison rather than, for example, the average over all other topics, means that a *k*-mer is only considered distinctive if it stands out against every other topic, not just most of them; this is a stricter criterion than the original formulation of [27], which uses a background reference distribution. A high score therefore indicates that a *k*-mer is reliably elevated in this topic in a way that cannot be attributed to any other topic. The threshold used to select driving *k*-mers is described in the Experimental Setup.

When running multiple model replicates, we apply a consensus filter: a *k*-mer is retained as a driving *k*-mer for a topic only if it appears in the driving list of more than 50% of the runs. This ensures that reported driving *k*-mers reflect stable, replicable signals rather than artifacts of a single model run. Among the *k*-mers that pass the consensus filter, the final ranking is determined by their distinctiveness score computed on the consensus-averaged topic distributions, so higher-ranked *k*-mers are both reproducible and strongly characteristic of the topic.

Importantly, the driving *k*-mers within each topic are both individually over-represented and tend to co-occur, forming a collective motif identification. The driving *k*-mers identified by KL-distinctiveness scoring characterize each topic by being the *k*-mers most specific to that topic relative to all others, in the comparison at hand. Therefore, all downstream analyses treat each topic as an indivisible motif family, computing distances and enrichment statistics jointly, over the entire set of driving *k*-mers.

## 3 Results

This section presents the experimental setup and the results of the topic modelling analysis.

### 3.1 Experimental Setup

This section outlines all experimental setups used in our experiments.

#### Genomic proxy

*R* = 10 independent genome proxies were generated for each genome, balancing representational breadth and computational cost. In line with the empirically determined optimal parameters of [20], we use a proxy length of 100 kbp which is the pseudo-concatenation of *n* = 10 random sub-fragments from the entire genome.

#### *k*-mer value selection

We evaluate two *k*-mer sizes: *k* = 6, and *k* = 9. The value *k* = 6 is suggested by prior work [20], where it was shown to be optimal for accurate taxonomic, as well as environment-type alignment-free classification. We additionally include the value *k* = 9, to explore the impact of capturing longer sequence motifs. Values of *k <* 6 were excluded since the canonical *k*-mer vocabulary at these sizes (32, 136, and 512 *k*-mers for *k* = 3, 4, 5 respectively) is too small, causing the *k*-mer frequency profiles across fragments to be nearly identical and providing insufficient variation for the model to identify distinct, reproducible topics. We also note that the value *k* = 3 was extensively studied in [13, 20], and corroborated with biology literature on codon and amino acid representation in extremophiles.

#### Topic modelling

The number of topics *T* was selected independently for each combination of domain, comparison, and *k*-mer size, using the `searchK` diagnostic [18], which evaluates held-out likelihood, semantic coherence, exclusivity, and residuals over a candidate range of *T* values. *T* was chosen at the point where held-out likelihood levels off while coherence and exclusivity remain acceptable. Table 2 summarizes the optimal number of topics, *T*, selected for each setting. The details of the `searchK` output are provided in Supplementary Materials, Section C. Ten independent STM initializations were used per configuration, which is sufficient to identify a stable consensus under the cosine-similarity alignment criterion while remaining computationally efficient.

**Table 2:**
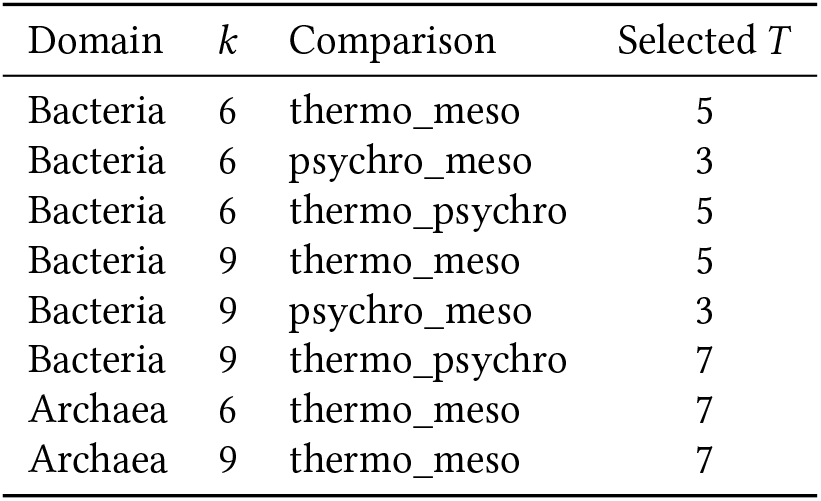
Optimal number of topics *T* selected via searchK for each computational experiment.

#### Driving *k*-mer selection

After fitting, each topic was assigned a stability score: the mean pairwise cosine similarity of its *k*-mer distribution across the ten runs. A stability score of 1 means the topic was recovered identically in every run; a score below 0.8 indicates the topic did not converge to a consistent solution and is therefore excluded from all downstream interpretation, its driving *k*-mers are not reported and it is not included in effect estimation. A threshold of 0.80 was chosen as a conservative cutoff, ensuring that only topics with a highly consistent *k*-mer structure across runs are carried forward into interpretation and effect estimation. For the remaining stable topics, *k*-mers were retained as driving *k*-mers if their KL distinctiveness score exceeded the within-topic mean by more than one standard deviation, a standard statistical threshold for identifying items that are meaningfully above average without requiring a hand-tuned cutoff.

### 3.2 Driving *k*-mers identified by STM

This section provides detailed results of the STM on our dataset.

#### 3.2.1 Co-occurring overrepresented hexamers in thermophiles

##### Bacterial species

The STM, thermo_meso comparison, (*T* =5) identified two topics, each comprising specific hexamers significantly overrepresented and co-occurring in bacterial thermophiles (*p <*0.05). The first topic (referred to as 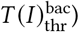 contains 30 hexamers identified as driving hexamers, and the second topic (referred to as 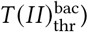 contains 26 hexamers identified as driving hexamers. Both topics contain exclusively G+C-rich hexamers, where a hexamer is termed to be *G+C-rich* if it contains at least four G/C bases. By this definition, 720 of the 2,080 canonical hexamers are G+C-rich, which implies that the STM-identified two topics are small, highly specific subsets of the G+C-rich hexamer pool. Specifically, the bacterial thermophile topics together comprise only 7.8% of all possible G+C-rich canonical hexamers.

Importantly, these motifs are recovered from a taxonomically diverse pool of bacterial thermophiles spanning 15 phyla, indicating that the identified motifs are not driven by a single lineage. The results for the thermo_psychro comparison are consistent with the results for thermo_meso comparison and are provided in Supplementary Materials, Section D.

The sequence logos for the hexamers of each of the two bacterial thermophile topics are shown in Figure 2. Each sequence logo summarizes the positional nucleotide composition of its topic’s hexamers: the height of a letter reflects how consistently that nucleotide appears in that position across all hexamers in the topic (measured in bits, where 2 bits indicates perfect conservation in that position.) A position where one letter dominates the others means that nucleotide is nearly fixed in that position across all hexamers in the topic; a position with several short letters of similar height means different nucleotides are tolerated at that position [24, 26]. For example, in the logo for 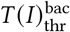 (Figure 2, (a)), the logo indicates a C- or G-stacked motif family, with C dominating most positions and G contributing consistently across the motif. Positions 5 and 6 show the strongest conservation, whereas earlier positions are more variable and show only weak A/T contributions.

**Figure 2:**
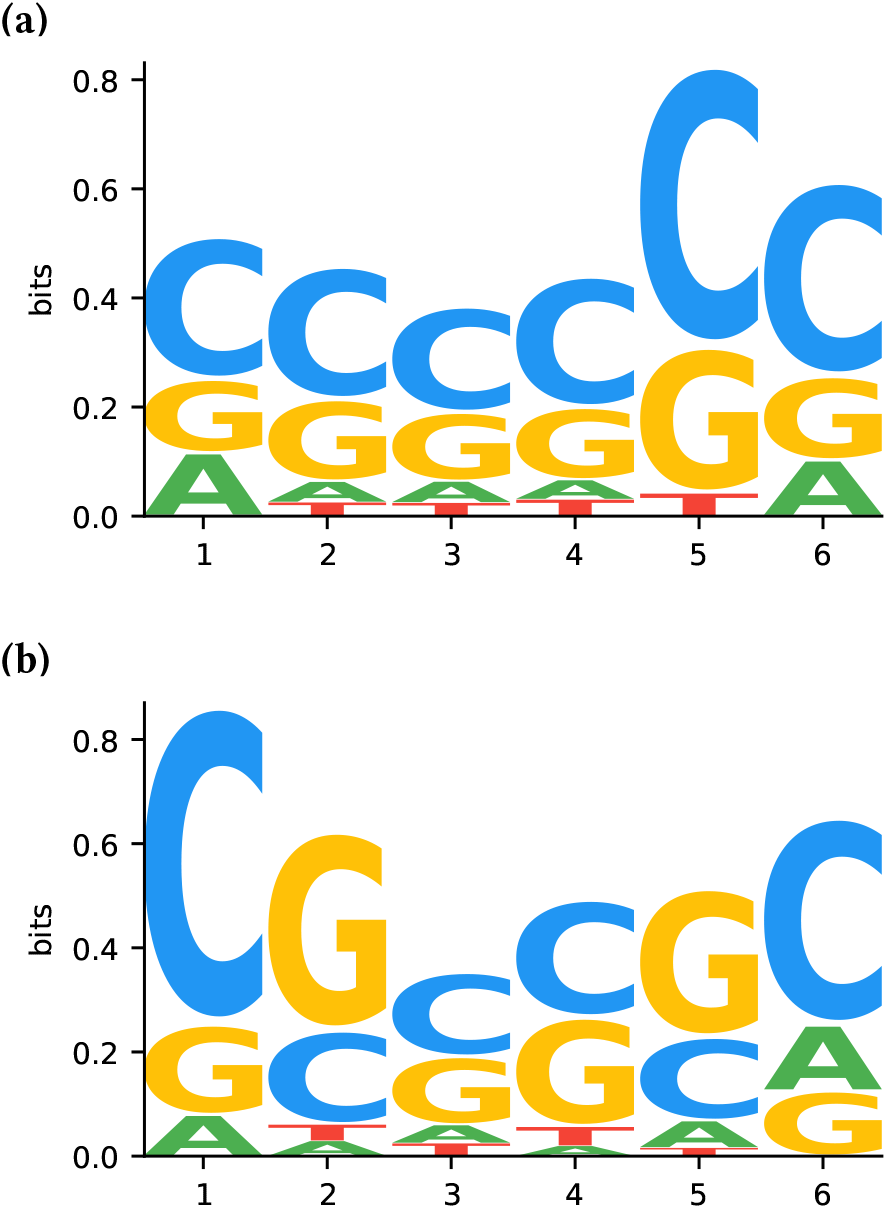
Thermophile-Enriched Sequence Motifs in Bacterial Species. Sequence logos showing hexamers overrepresented and co-occurring in thermophilic bacterial topics. (**a**) Driving hexamers identified in topic 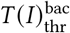, (**b**) driving hexamers identified in topic 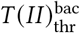.

##### Archaeal species

The STM, thermo_meso comparison, (*T* =7) identified one topic, comprising specific hexamers significantly overrepresented and co-occurring in archaeal thermophiles (*p <*0.05). This topic (referred to as 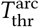) has 16 driving hexamers, unified by a core `CTAG` tetranucleotide motif, with 7 out of 16 hexamers (43.8%) meeting the G+C-rich criterion (e.g., CTAGCC, GCTAGC, CGCTAG, CCCTAG). This indicates that the archaeal thermophilic signal is driven by the conserved `CTAG` core rather than only G+C enrichment. Note that these 7 hexamers represent only 0.97% of all 720 possible canonical G+C-rich hexamers. This partially parallels the bacterial signal, in that the thermophile-associated hexamers are G+C-biased in both cases, yet the archaeal pattern is more centered around a single motif. Importantly, this signature is recovered from a taxonomically diverse pool of archaeal thermophiles spanning 6 phyla, indicating that the `CTAG`-centred signal is not confined to a single taxonomic lineage. The sequence logo for this topic, 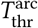, is shown in Figure 3.

**Figure 3:**
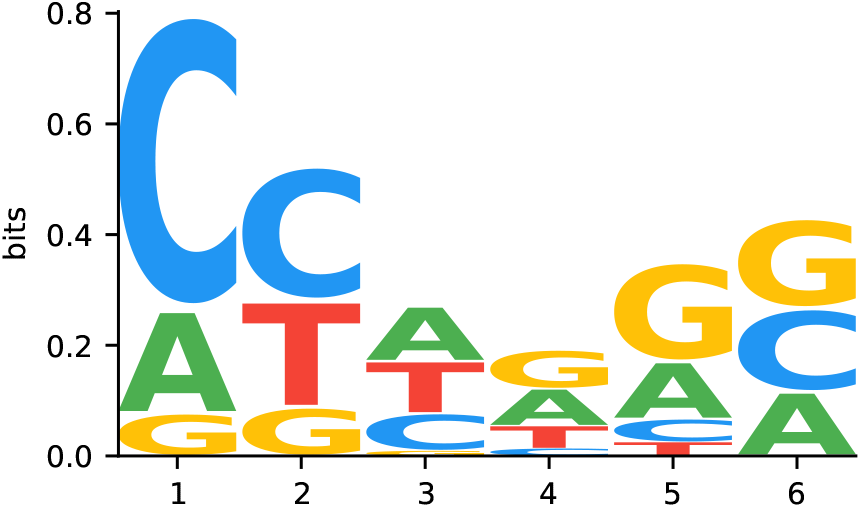
Thermophile-Enriched Sequence Motifs in Archaeal Species. Sequence logos showing hexamers overrepresented and co-occurring in thermophilic archaeal topics, with driving *k*-mers identified in topic 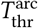.

Table 3 summarizes the overrepresented and co-occurring hexamers for thermophilic bacteria and archaea across all comparisons. In bacterial thermophiles, two distinct but related classes emerge: a class of C- or G-stacked motifs, and a class of CG-periodic motifs. In archaeal thermophiles, the identified motifs define a coherent `CTAG`-centred family, highlighting that different domains may realize the same selective pressure through distinct sequence-level solutions.

**Table 3:**
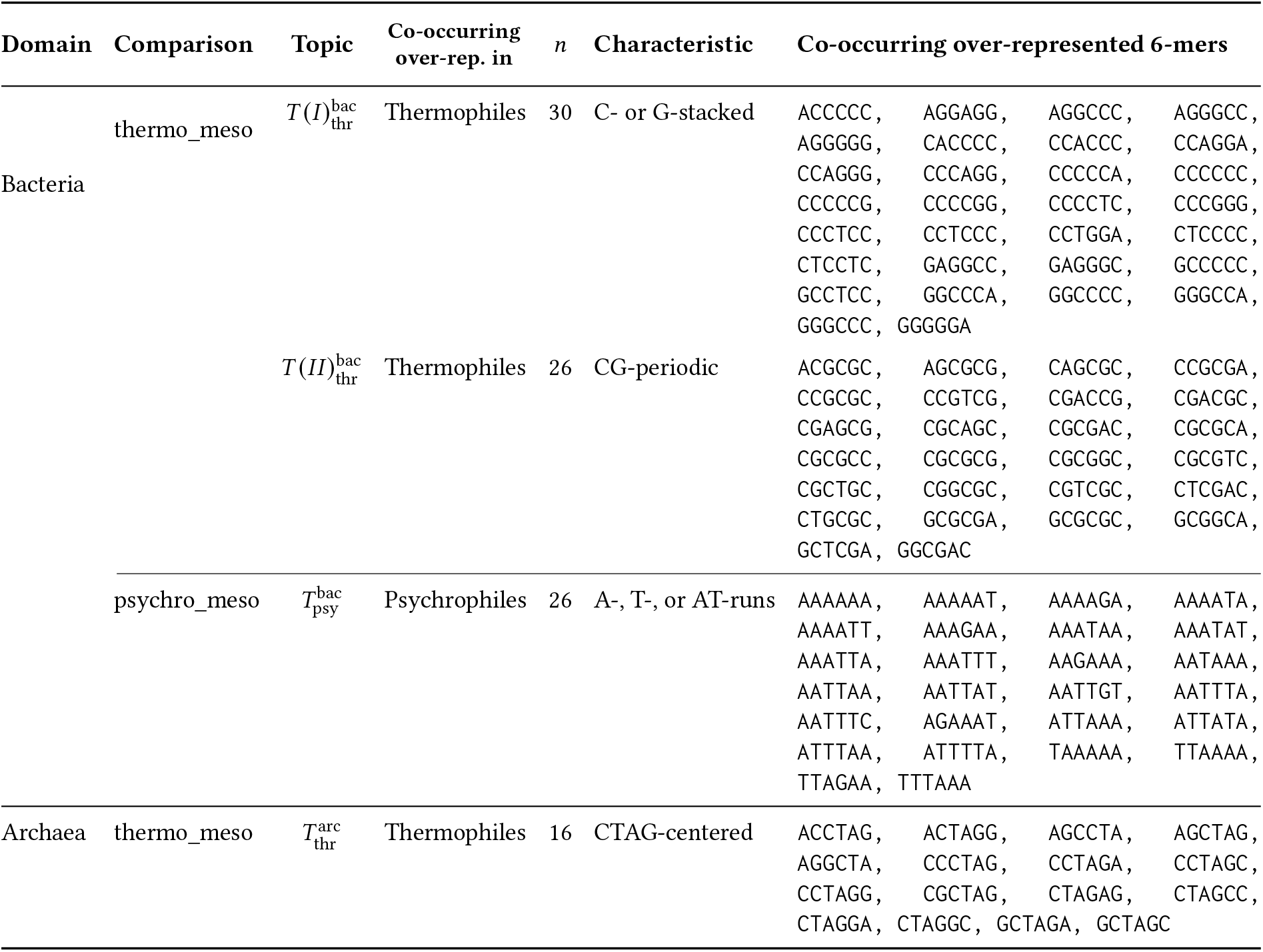
Co-occurring over-represented canonical 6-mers per STM topic, grouped by domain and comparison.

Importantly, these motif-family signals are not detectable at lower *k* values (*k <* 6), where the canonical *k*-mer vocabulary is too small for STM to identify distinct and reproducible topics. This observation is consistent with previous studies showing that genome representations based solely on canonical 1-mer frequencies (equivalent to overall G+C content) possess limited discriminatory power for environment-type classification in extremophiles. Using either whole-genome representatives or randomly sampled genome proxies, classification accuracies based on *k* = 1 profiles remained only modestly above random expectations [13, 20]. Together, these results indicate that the thermal-adaptation signal identified here does not arise primarily from overall nucleotide composition, but from highly selective sequence architectures represented by specific co-occurring motif families.

#### 3.2.2 Co-occurring overrepresented hexamers in psychrophiles

##### Bacterial species

The STM psychro_meso comparison (*T* =3) identified one topic comprising specific hexamers significantly overrepresented and co-occurring in bacterial psychrophiles (*p* <0.05). This topic (referred to as 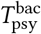), with 26 driving hexamers, contains exclusively A+T-rich motifs (e.g., AAAAAA, TTAAAA, AAAATT, ATTTTA), consistent with the lower GC content characteristic of cold-adapted genomes. Note also that this topic comprises only 3.6% of the 720 possible canonical A+T-rich hexamers. The same pattern is found in the thermo_psychro comparison (*T* =5), and the results are provided in Supplementary Materials, Section D. Notably, these driving hexamers are recovered from a dataset containing bacterial psychrophiles spanning 4 phyla, indicating that this signal is not taxonomic lineage-specific. Sequence logos of the overrepresented and co-occurring hexamers in bacterial psychrophiles are shown in Figure 4. Table 3 summarizes the overrepresented and co-occurring hexamers for psychrophilic bacteria across both comparisons. In contrast to thermophiles, bacterial psychrophile-associated hexamers are low-complexity A-, T-, or AT-runs.

**Figure 4:**
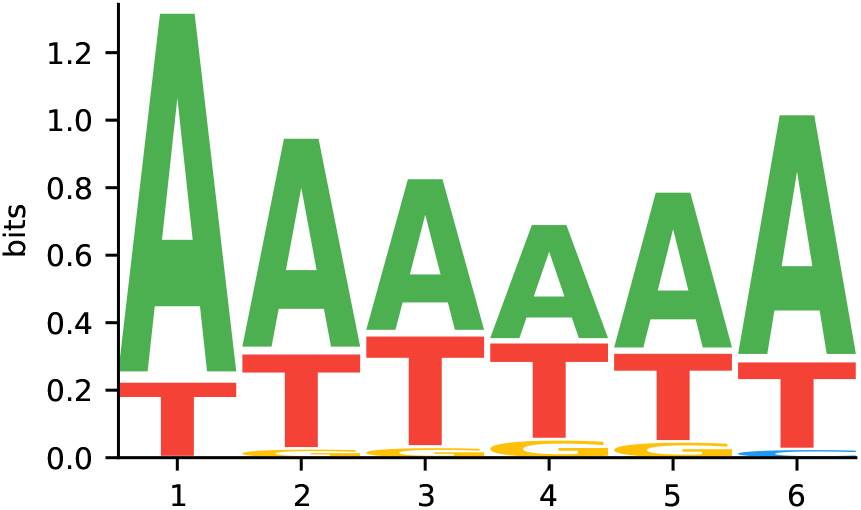
Psychrophile-Enriched Sequence Motifs in Bacterial Species. Sequence logos showing hexamers overrepresented and co-occurring in psychrophilic bacterial topics. (**a**) Driving hexamers identified in 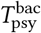.

##### Archaeal species

The dataset contains only 8 psychrophile archaeal genomes, which is insufficient for reliable STM application; these species are instead analyzed separately and the results are provided in Supplementary Materials, Section E.

#### 3.2.3 Overrepresentation of 9-mers in thermophiles

##### Bacterial species

The bacterial thermophile signal at *k*=9 is fully consistent with *k*=6. All 30 bacterial thermophile driving 9-mers identified by the STM (thermo_meso, *T* =5) are G+C-rich (defined as ≥ 5 G/C bases in 9-mers), covering only 0.046% of all 65,536 possible canonical G+C-rich 9-mers. Furthermore, 90% of these 9-mers contain at least one overrepresented hexamer as a substring, confirming that the 9-mers are extensions of the same core hexamer motifs. Full 9-mer lists and sequence logos are provided in Supplementary Materials, Sections F and G, respectively.

##### Archaeal species

Only four 9-mers passed the topic stability filters, forming a weak motif signal. Full 9-mer lists and sequence logos are provided in Supplementary Materials, Sections F and G, respectively.

#### 3.2.4 Overrepresentation of 9-mers in psychrophiles

##### Bacterial species

The bacterial psychrophile signal at *k*=9 is consistent with overrepresented hexamers. All 21 bacterial psychrophile driving 9-mers (psychro_meso, *T* =3) are A+T-rich (≥ 5 A/T bases), and all of them contain at least one overrepresented hexamer as a substring. These 9-mers cover only 0.032% of all 65,536 possible canonical A+T-rich 9-mers. Full 9-mer lists and sequence logos are provided in Supplementary Materials, Sections F and G, respectively.

### 3.3 Cross-domain hexamer convergence in two thermophilic bacterium–archaeon pairs

The STM analysis identified distinct thermophile-associated motif families in bacteria and archaea: C- or G-stacked and CG-periodic motif families in bacterial thermophiles, and a CTAG-centred motif family in archaeal thermophiles. Although these results suggest domain-specific sequence-level responses to high temperature, they do not exclude the possibility that particular bacterial and archaeal species may converge toward similar motif usage under shared environmental pressures. We therefore explored the hypothesis that some divergent thermophilic bacterium–archaeon pairs may have similar frequencies of the thermophile-associated motifs identified in either domain.

To address this question, we examined two well-characterised thermophilic bacterium–archaeon pairs previously identified in [20] as carrying highly similar genomic signatures despite maximal taxonomic divergence: Pair 1, comprising the polyextremophiles *Thermoanaerobacterium thermosaccharolyticum* and *Caldisphaera lagunensis*, thriving at high temperature and low pH, and Pair 2, comprising *Thermotoga petrophila* and *Geoglobus acetivorans*, both hyperthermophiles isolated from high-temperature hydrothermal environments. For each pair, we computed the normalized Manhattan distance and Jensen–Shannon (JS) distance between the canonical hexamer frequency profiles of the two species, restricted to three sets of thermophile-associated hexamers identified by STM: the two bacterial thermophile topics, 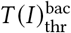 and 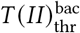, and the archaeal thermophile topic, 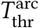. Computing distances separately for each topic’s driving hexamers allowed us to test whether the members of each pair were similar with respect to the bacterial thermophile motif families, the archaeal thermophile motif family, or both. As a baseline, we computed the same distances among thermophilic genomes within each domain, providing a reference for the expected within-domain hexamer distance under the same driving hexamer sets.

As shown in Figure 5, in both pairs, the hexamer distance between the bacterium and the archaeon was substantially smaller than the mean distance observed between thermophilic genomes within the same domain. Thus, in these two specific cases, the paired bacterial and archaeal species are more similar to each other in their usage of thermophile-associated hexamers than expected from typical same-domain thermophile comparisons. This suggests that, although the dominant thermophile-associated motif families differ between bacteria and archaea overall, individual species from the two domains can converge toward similar usage of both bacterial and archaeal thermophile-associated motif sets under shared high-temperature conditions. Further investigations, of larger and more diverse extremophile datasets, are needed to elucidate the mechanisms behind such cross-domain convergence in thermophile-associated motif usage.

**Figure 5:**
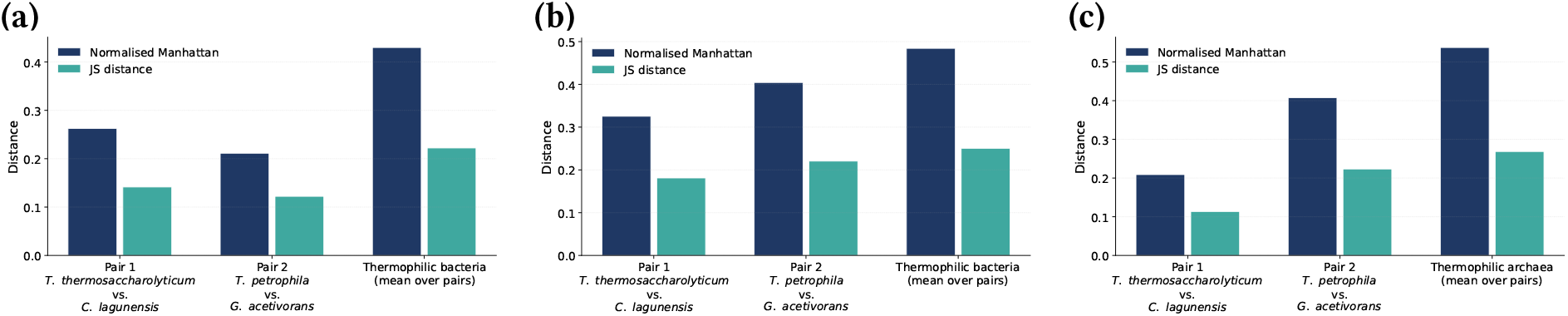
Cross-Domain Driving Hexamer Distance in Two Thermophilic Bacterium–Archaeon Pairs. The normalised Manhattan distance and Jensen–Shannon (JS) distance between the canonical hexamer frequency profiles of two cross-domain thermophilic pairs and the within-domain thermophile baseline, were computed over the driving hexamers of each STM topic. (**a**) 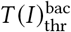: C- or G-stacked motifs. (**b**) 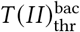: CG-periodic motifs. (**c**) 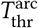: CTAG-centred motifs. **Pair 1**: *T. thermosaccharolyticum* (bacterium) and *C. lagunensis* (archaeon), both polyextremophiles thriving at high temperature and low pH. **Pair 2**: *T. petrophila* (bacterium) and *G. acetivorans* (archaeon), both isolated from high-temperature hydrothermal environments. The within-domain baseline represents the mean pairwise distance between thermophilic genomes within the same domain. Both pairs show lower cross-domain distances than the within-domain baseline across all driving hexamer sets, suggesting convergent genome-wide sequence composition under shared thermal selective pressure.

## 4 Discussion

Our analysis reveals a strong, highly structured, and reproducible signal of thermal adaptation in the form of genome-wide distribution of specific short DNA sequences. The motif sets identified by the STM comprise four distinct topic families, each recovered consistently across comparison settings and extending coherently to the 9-mer scale: 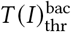 (a set of C- or G-stacked hexamers) and 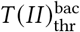 (a set of CG-periodic hexamers), both over-represented and co-occurring in bacterial thermophiles; 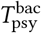 (a set of A-, T-, or AT-run low-complexity hexamers) overrepresented and co-occurring in bacterial psychrophiles; and 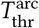 (a set of hexamers unified by a conserved `CTAG` core) overrepresented and co-occurring in archaeal thermophiles. Given that previous studies have shown DNA structural and mechanical properties to depend on local sequence context, including at the hexanucleotide scale [25, 11], we now interpret each STM-identified hexamer family in relation to known sequence-dependent DNA physical properties.

Thermophile-associated bacterial motifs exhibit two related but distinct patterns.

The first topic, 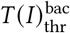, is dominated by C- or G-stacked sequence motifs and, notably, the TpA dinucleotide step is entirely absent. Previous studies found the TpA dinucleotide to be the most conformationally flexible [11]. Thus, the absence of TpA in the hexamers of 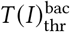 suggests depletion of locally flexible sequence contexts. We also note that the G- or C-stacking composition of the 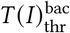 hexamers is mechanically relevant. Indeed, previous studies showed that base-stacking is the main stabilizing factor in the DNA double helix, and that base-pair steps involving G and C contribute more strongly to stacking stability than those involving A and T [29].

A closer analysis of the first topic, 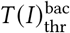, reveals that 17 of the 30 hexamers contain at least one GG dinucleotide, reflecting a bias toward G-stacked sequence contexts. We note that consecutive guanine bases have been known to form particularly stable stacking interactions, conferring additional local rigidity to the duplex [11]. Moreover, it has been observed that G-rich sequences capable of forming G-quadruplex structures are enriched in thermophilic bacterial genomes [4]. Lastly, a recent study showed that G-quadruplex potential (specifically within 16S rRNA gene regions) correlates positively with high optimal growth temperature across prokaryotes, whereas genomic G+C content does not [10]. These biological observations parallel our finding that specific G-stacked motif classes, rather than G+C content per se, carry the thermal adaptation signal. Together, these findings raise the possibility that the enrichment of specific G-stacked motifs partly reflects selection for G-quadruplex-forming potential in functionally important genomic regions.

The 26 driving hexamers of the second thermophilic bacterial topic, 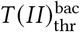, include motifs such as CGCGCG, CGCGCC, and `CGCGAC`, and are characterized by a pronounced CG-periodic structure and a complete absence of TpA motifs. The interpretation of the absence of TpA motifs parallels that of the first thermophilic bacterial topic. The CG-periodic structure is directly related to repeated CpG/GpC dinucleotide steps. Note that CpG-rich sequences have been previously identified as relatively rigid DNA regions [11], and their enrichment may therefore reflect selection for reduced local flexibility at high temperature. A complementary interpretation is that the periodic arrangement of CpG/GpC steps may influence how DNA responds to torsional stress [17], which could be relevant during replication and transcription at high temperature. Whether this mechanism underlies the enrichment of 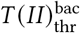 remains an open question requiring further experimental investigation.

Psychrophile-associated bacterial motifs exhibit patterns contrasting with those of thermophilic bacteria. Indeed, all 26 driving hexamers of 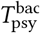, e.g., AAAAAA, TTAAAA, AAAATA, and `TATAAA`, are A-, T-, or AT-runs. As previously noted, the TpA dinucleotide is among the most conformationally flexible dinucleotide steps [11, 1], and alternating A+T-rich motifs contain repeated flexible steps. In addition, A+T-rich sequence contexts generally have weaker stacking contributions to duplex stability than G+C-rich contexts [29]. Thus, the enrichment of A-, T-, and AT-run motifs in psychrophilic genomes is consistent with a shift toward sequence contexts with lower local duplex stability and potentially greater local deformability. Such properties may help maintain local DNA flexibility at low temperatures, where molecular motion is reduced.

In contrast to bacteria, the hexamers in topic 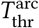 are unified by a core `CTAG` tetranucleotide rather than by C- and G-stacked or CG-periodic motifs. This observation differs from earlier reports of `CTAG` depletion, which were derived primarily from analyses of haloarchaeal genomes [5, 15] and from two thermophilic methanogenic archaea, *M. jannaschii* and *M. thermoautotrophicum* [8]. By contrast, our dataset comprises 229 archaeal genomes, including 137 thermophiles spanning a broad taxonomic range. Within this larger collection, `CTAG`-containing hexamers are consistently enriched and co-occurring in thermophilic archaeal genomic signatures, and a separate motif-frequency analysis shows that `CTAG` occurs 3.41-fold more frequently in archaeal thermophiles than in archaeal mesophiles. These results suggest that enrichment of `CTAG`-centred motifs may be a more widespread feature of archaeal thermal adaptation than previously recognized. The finding is particularly intriguing because `CTAG` contains the conformationally flexible TpA dinucleotide, which in our analyses is associated with psychrophile-enriched motifs in bacteria. This contrast suggests that thermal adaptation may involve domain-specific modes of DNA sequence organization rather than a universal shift toward maximal duplex rigidity.

The two examples of cross-domain hexamer profile convergence provide a more nuanced view of the relationship between phylogeny and thermal adaptation. On the one hand, at the global level, bacteria and archaea appear to use distinct thermophile-associated motif families, suggesting domain-specific sequence-level responses to high temperature. On the other hand, the two thermophilic bacterium–archaeon pairs examined here show unusually similar frequencies across both bacterial and archaeal thermophile-associated motif sets. This indicates that, in these two specific case studies, the bacterial and archaeal genomes have become more similar to each other in their representation of thermophile-associated hexamers, despite belonging to different domains of life. This finding supports the idea that shared extreme thermal pressure can, in some cases, partially override the expected taxonomic signature in genome-wide motif usage.

## 5 Conclusions and Future Work

Taken together, our results show that genome-wide distributions of short DNA sequences carry a robust and selective signal of thermal adaptation that is not explained by simple shifts in G+C or A+T content alone, but instead by the repeated enrichment of specific co-occurring overrepresented hexamer motif families. Remarkably, among the 720 possible G+C-rich hexamers, only 56 are associated with bacterial thermophiles, while among the 720 possible A+T-rich hexamers, only 26 are associated with bacterial psychrophiles. Thus, thermal adaptation is associated not with broad compositional trends, but with a surprisingly small and reproducible subset of sequence motifs. The thermophile- and psychrophile-associated motif classes identified here are consistent with distinct regimes of sequence-dependent DNA physical behaviour and suggest that adaptation to temperature is reflected in highly specific genome-wide sequence architectures. Together, these findings provide a mechanistic interpretation for how genome-wide sequence organization may encode adaptation to temperature beyond phylogenetic history.

Future work should focus on establishing a more direct quantitative connection between the motif families identified here and sequence-dependent DNA physical properties. In particular, it will be important to determine whether thermophile- and psychrophile-associated motifs occupy distinct regions of sequence-property space defined by experimentally measured or computationally predicted parameters such as base-stacking energetics, conformational flexibility, and DNA shape. Such analyses would provide a direct test of the mechanistic hypothesis suggested by the present results. In addition, the distinct CTAG-centred motif family identified in thermophilic archaea warrants further investigation, as it may represent an alternative sequence-level solution to thermal adaptation. More broadly, extending this framework to other environmental variables and larger microbial datasets may help reveal whether genome-wide sequence organization encodes a general physical layer of adaptive information beyond temperature alone.

## 6 Acknowledgments

Author Contributions: L.K. conceived the study and interpreted the results; M.S. developed the methodology and performed the analyses; both authors wrote the manuscript.

