## Supplementary Materials for "Beyond phylogeny: Genome-wide DNA sequence patterns suggest DNA physical properties associated with thermal adaptation in extremophile microbes"

#### A Dataset Details

Figure S1 and Figure S2 provide the Sunburst diagrams for the bacterial and archaeal domains of the Temperature Dataset, illustrating taxonomic diversity from phylum to genus level.

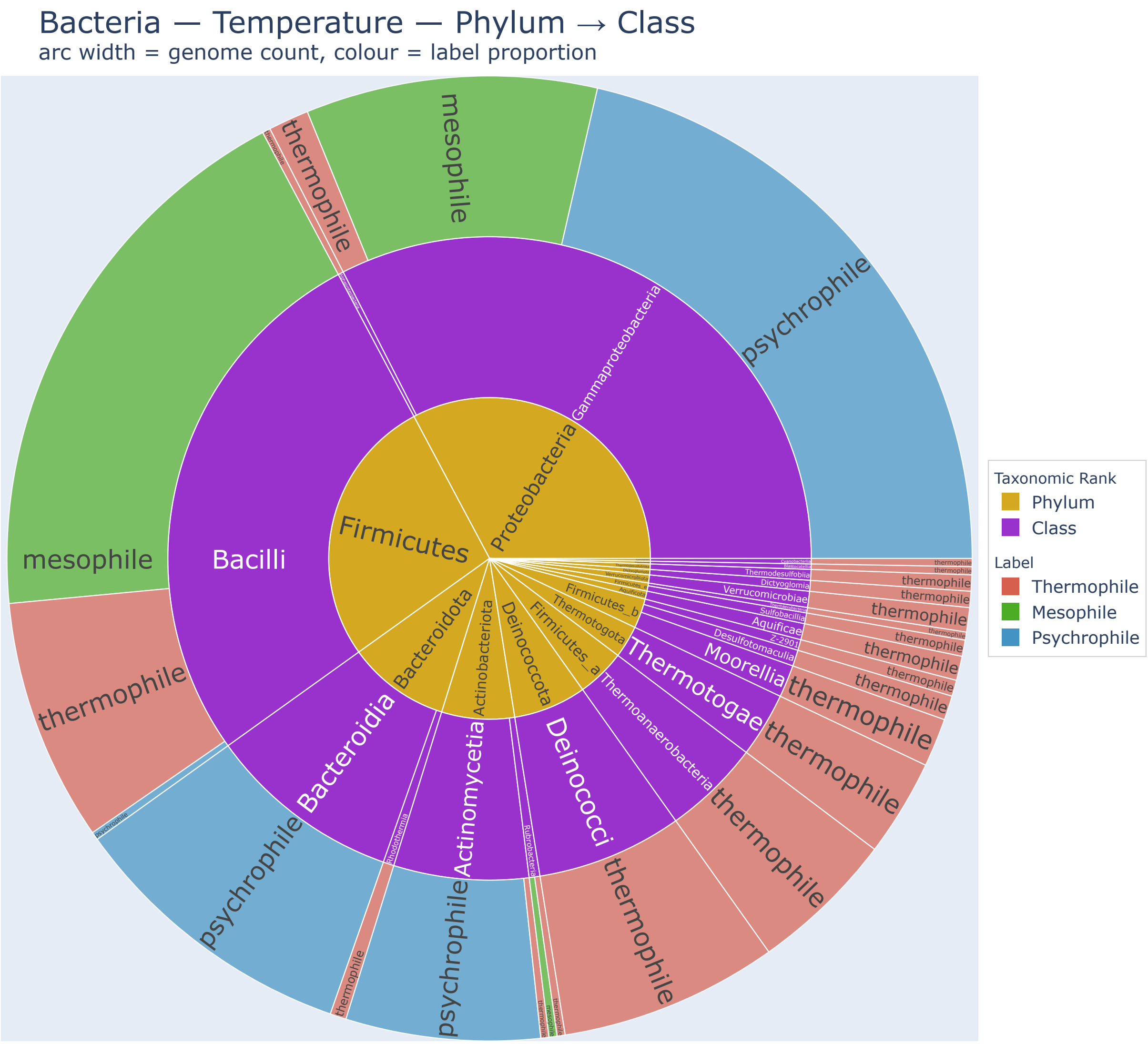


Figure S1: Sunburst diagram for the phylum and class taxonomic level of all the bacterial species in the Temperature dataset.

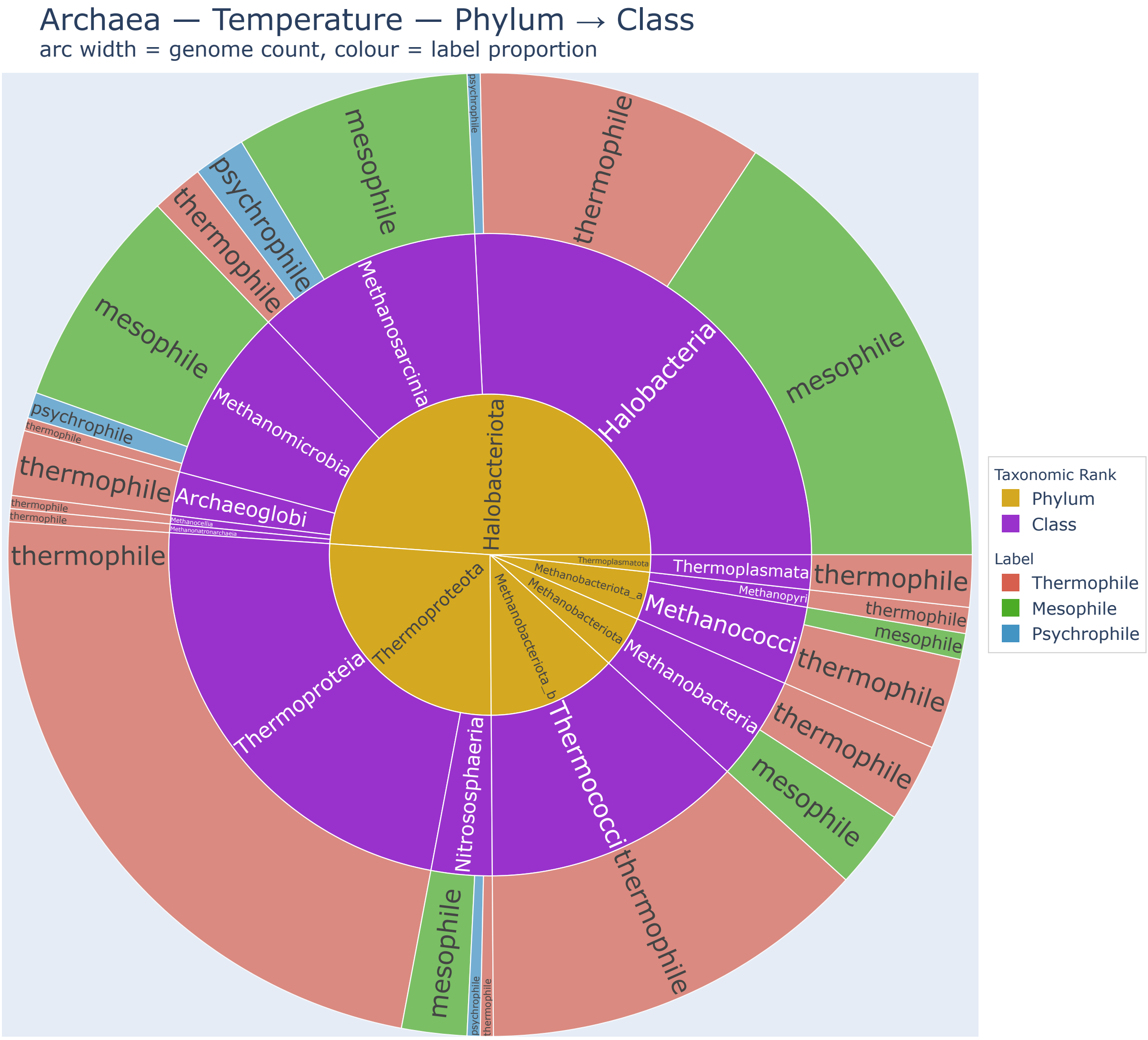


Figure S2: Sunburst diagram for the phylum and class taxonomic level of all the archaeal species in the Temperature dataset.

### B Technical Details of the STM Prevalence Formula

#### What the prevalence formula does

In STM, each document (here a genome proxy) is described by a mixture of topics, where each topic is a probability distribution over all $k$-mers. The *prevalence formula* tells the model which external variables may shift how much a genome uses each topic. It does not change the topics themselves; it only allows topic usage to vary systematically between groups.

Our prevalence formula is:

```
~ temperature_label + Genus
```

**Temperature label.** A categorical variable indicating whether a genome belongs to Thermophiles, Psychrophiles, or Mesophiles (reference level). The model estimates, for each topic, how much more or less that topic is used in thermophiles or psychrophiles compared with mesophiles.

**Genus covariate.** Included to account for phylogenetic relatedness: closely related genomes tend to share $k$-mer composition regardless of temperature adaptation. By including Genus, the model controls for this background signal so that the temperature effect reflects adaptation rather than ancestry. Genera with fewer than two representatives are excluded before fitting.

##### How topics are linked to temperature groups

After fitting the STM, genome-level topic proportions (averaged across $R = 10$ genome proxies per genome) are centred log-ratio (CLR) transformed to remove the compositional constraint (topic proportions must sum to 1). For each topic $k$ we then fit a linear regression:

$$\tilde{\theta}_{ik} = \alpha + \beta\,\text{label}_i + \gamma\,\text{Genus}_i + \varepsilon_i$$

A positive coefficient $\beta$ means the topic is more prevalent in the target group (e.g., thermophiles) compared with the mesophile baseline. $p$-values are Benjamini–Hochberg corrected across all topic–label combinations; topics with corrected $p < 0.05$ and $\beta > 0$ are reported as significantly enriched in the target group.

#### C `searchK` Diagnostic Output

##### How we chose the number of topics

The number of topics $T$ is the main hyperparameter of an STM. To choose it, we used the `searchK` function from the `stm` R package, which fits models with different values of $T$ and evaluates each on four metrics: held-out likelihood (how well the model predicts unseen data), semantic coherence (whether the top words of each topic frequently co-occur), exclusivity (whether top words are specific to one topic rather than shared across many), and residuals. For each configuration (domain, $k$-mer size, and comparison group) we examined the resulting diagnostic plots and selected the value of $T$ at which held-out likelihood levels off while semantic coherence and exclusivity remain acceptable—i.e., the "elbow" of the held-out likelihood curve, beyond which adding more topics provides diminishing returns. The selected values are shown in Table S1, and the corresponding diagnostic plots are shown in Figures S3, S4, and S5.

Table S1: Optimal number of topics $T$ selected via `searchK` for each domain, k-mer size, and comparison setting.

| Domain | $k$ | Comparison | Selected $T$ |
|---|---|---|---|
| Bacteria | 6 | thermo-meso | 5 |
| Bacteria | 6 | psychro-meso | 3 |
| Bacteria | 6 | thermo-psychro | 5 |
| Bacteria | 9 | thermo-meso | 5 |
| Bacteria | 9 | psychro-meso | 3 |
| Bacteria | 9 | thermo-psychro | 7 |
| Archaea | 6 | thermo-meso | 7 |
| Archaea | 9 | thermo-meso | 7 |

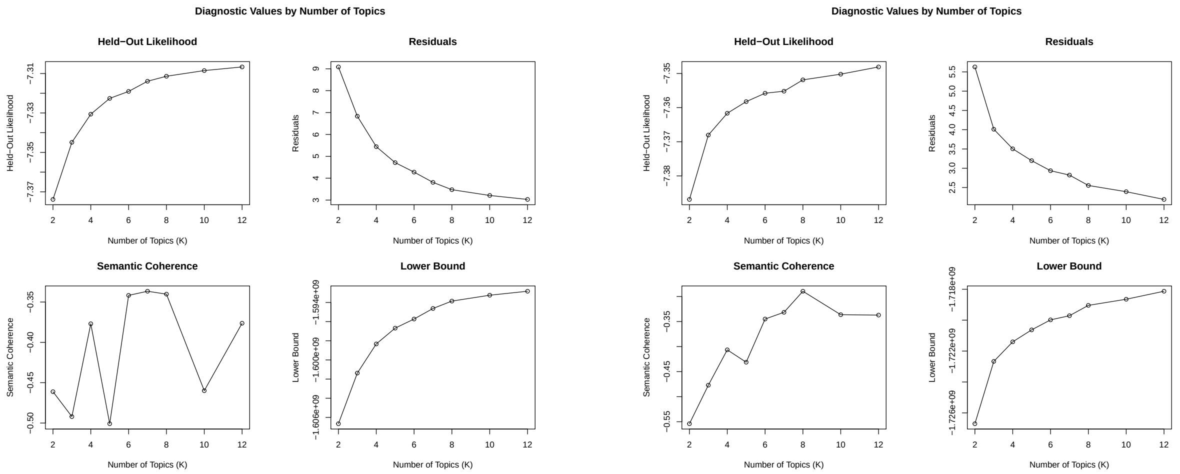


Figure S3: `searchK` diagnostics for bacteria, $k = 6$. Left: thermo-meso ($T = 5$ selected). Right: psychro-meso ($T = 3$ selected).

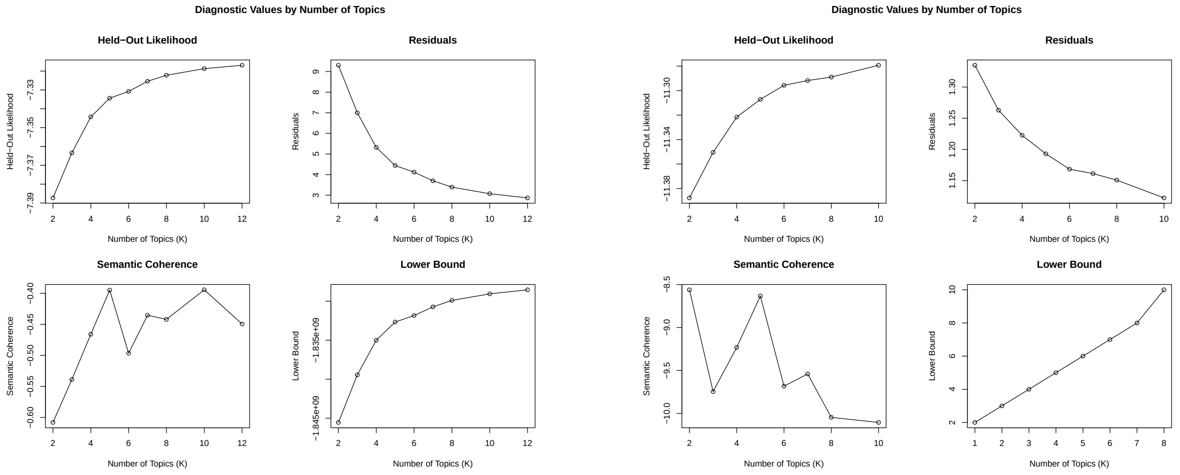


Figure S4: `searchK` diagnostics for bacteria, thermo-psychro. Left: $k = 6$ ($T = 5$ selected). Right: $k = 9$ ($T = 7$ selected).

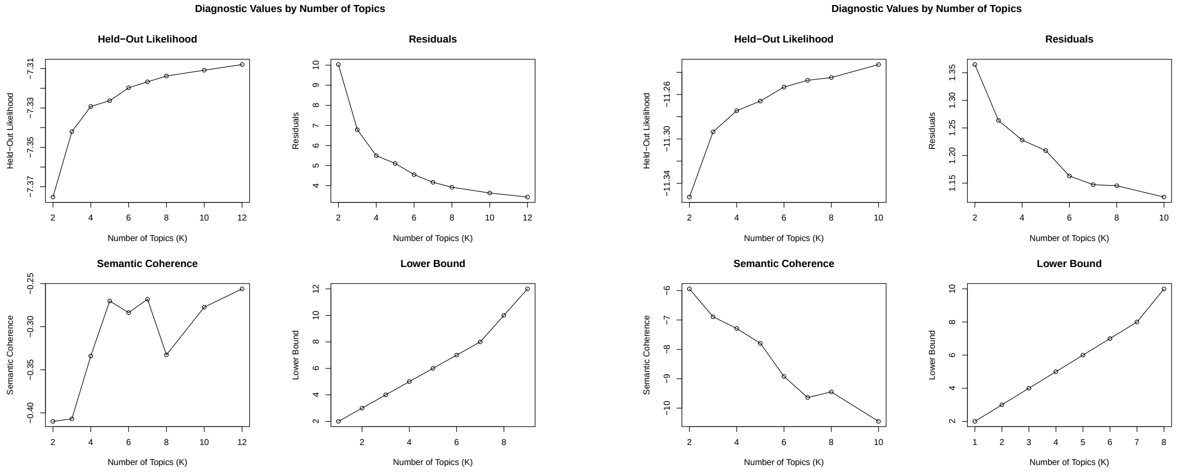


Figure S5: `searchK` diagnostics for archaea, thermo-meso. Left: $k = 6$ ($T = 7$ selected). Right: $k = 9$ ($T = 7$ selected).

### D Results of Psychro-Thermo Comparison in Bacteria

We performed the STM analysis for the thermo-psychro comparison. The same GC-rich signature is recovered in this setting, where one significant topic contains 27 CG-rich driving hexamers, mirroring the thermophile signal observed previously. As in the psychro–meso comparison, the AT-rich psychrophile signal was also recovered: Topic 4 comprises 19 AT-rich driving hexamers (e.g., `AAAAAA`, `TTAAAA`, `AAAATT`). The two psychrophile-enriched topics identified in the psychro-meso and thermo-psychro comparisons share 17 hexamers (65% and 89% of each topic, respectively), confirming that the AT-rich signature is robust and independent of the choice of comparison group.

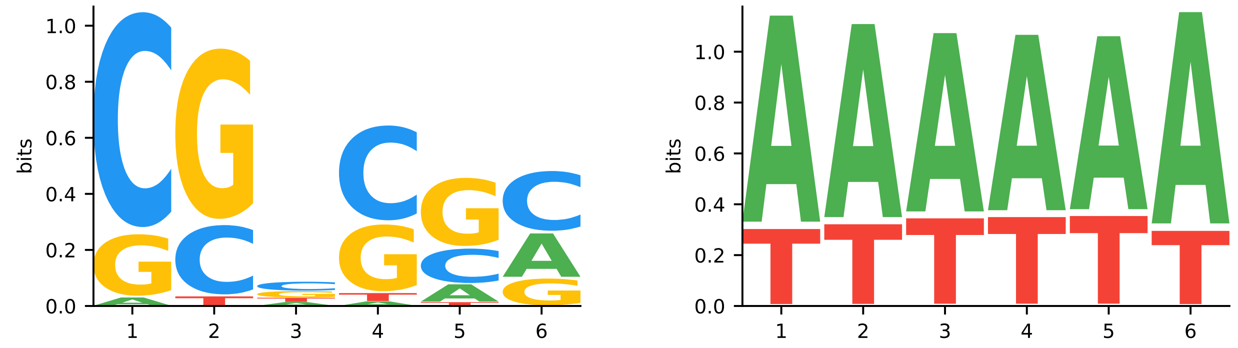


Figure S6: Sequence logos of overrepresented topics in the thermo-psychro comparison in bacteria. Left: thermophile-overrepresented topic. Right: psychrophile-overrepresented AT-rich topic.

Table S2: Driving hexamers identified in the *thermo-psychro* comparison ($T = 5$).

| **Group** | $n$ | **Driving hexamers** |
|---|---|---|
| Thermophiles | 27 | `AGCGCG, CCGACG, CCGCGA, CCGCGC, CCGTCG, CGACCG, CGACGA, CGACGC, CGAGCG, CGCCGA, CGCGAC, CGCGCA, CGCGCC, CGCGCG, CGCGGC, CGCGTC, CGGCGA, CGGCGC, CGTCGA, CGTCGC, CTCGAC, GCCGAC, GCGCGA, GCGCGC, GCTCGA, GGCGAC, GGTCGA` |
| Psychrophiles | 19 | `AAAAAA, AAAAAT, AAAATA, AAAATT, AAATAA, AAATAT, AAATTA, AAATTT, AATAAA, AATTAA, AATTAT, AATTTA, ATAAAA, ATTATA, ATTTTA, TAAAAA, TATAAA, TTAAAA, TTTAAA` |

### E Cross-Domain Analysis: Archaeal Psychrophiles

To investigate whether the AT-rich hexamer motifs identified in bacterial psychrophiles are detectable in archaeal psychrophiles, we projected archaeal genome proxies into the pre-trained bacterial psychrophile topic space using `fitNewDocuments`, which assigns topic weights based solely on the topic-word distributions learned from bacteria. This was necessary because the archaeal dataset contains only 8 psychrophile genomes, insufficient for reliable independent STM fitting.

Among the 8 archaeal psychrophile genomes, 5 placed dominant weight on the AT-rich, low-complexity hexamers enriched in bacterial psychrophiles ($T_{\text{psy}}^{\text{bac}}$), with combined loading between 0.78 and 0.82. One genome placed nearly all its weight ($\approx$ 0.99) on non-AT-rich mesophile-associated hexamers, while the remaining two showed intermediate profiles. Overall, the majority of archaeal psychrophile genomes display the same AT-rich motif bias identified in bacterial psychrophiles, suggesting a shared, domain-spanning genomic response to cold adaptation.

### F Full Driving 6-mer and 9-mer Lists per Topic

Tables S3 and S4 list the driving hexamers and 9-mers for each topic significantly enriched in the target temperature group, across all domain–comparison configurations.

Table S3: Over-represented canonical 6-mers per STM topic. Where multiple topics within the same comparison are enriched in the same temperature group, their driving $k$-mers are combined into a single row ($n$ = total count).

| **Domain** | **Comparison** | **Over-rep. in** | $n$ | **Over-represented 6-mers** |
|---|---|---|---|---|
| Bacteria | thermo-meso | Thermophiles | 56 | ACGCGC, AGCGCG, CAGCGC, CCGCGA, CCGCGC, CCGTCG, CGACCG, CGACGC, CGAGCG, CGCAGC, CGCGAC, CGCGCA, CGCGCC, CGCGCG, CGCGGC, CGCGTC, CGCTGC, CGGCGC, CGTCGC, CTCGAC, CTGCGC, GCGCGA, GCGCGC, GCGGCA, GCTCGA, GGCGAC, ACCCCC, AGGAGG, AGGCCC, AGGGCC, AGGGGG, CACCCC, CCACCC, CCAGGA, CCAGGG, CCCAGG, CCCCCA, CCCCCC, CCCCCG, CCCCGG, CCCCTC, CCCGGG, CCCTCC, CCTCCC, CCTGGA, CTCCCC, CTCCTC, GAGGCC, GAGGGC, GCCCCC, GCCTCC, GGCCCA, GGCCCC, GGGCCA, GGGCCC, GGGGGA |
| Bacteria | psychro-meso | Psychrophiles | 26 | AAAAAA, AAAAAT, AAAAGA, AAAATA, AAAATT, AAAGAA, AAATAA, AAATAT, AAATTA, AAATTT, AAGAAA, AATAAA, AATTAA, AATTAT, AATTGT, AATTTA, AATTTC, AGAAAT, ATTAAA, ATTATA, ATTTAA, ATTTTA, TAAAAA, TTAAAA, TTAGAA, TTTAAA |
| Bacteria | thermo-psychro | Thermophiles | 27 | AGCGCG, CCGACG, CCGCGA, CCGCGC, CCGTCG, CGACCG, CGACGA, CGACGC, CGAGCG, CGCCGA, CGCGAC, CGCGCA, CGCGCC, CGCGCG, CGCGGC, CGCGTC, CGGCGA, CGGCGC, CGTCGA, CGTCGC, CTCGAC, GCCGAC, GCGCGA, GCGCGC, GCTCGA, GGCGAC, GGTCGA |
| | | Psychrophiles | 19 | AAAAAA, AAAAAT, AAAATA, AAAATT, AAATAA, AAATAT, AAATTA, AAATTT, AATAAA, AATTAA, AATTAT, AATTTA, ATAAAA, ATTATA, ATTTTA, TAAAAA, TATAAA, TTAAAA, TTTAAA |
| Archaea | thermo-meso | Thermophiles | 16 | ACCTAG, ACTAGG, AGCCTA, AGCTAG, AGGCTA, CCCTAG, CCTAGA, CCTAGC, CCTAGG, CGCTAG, CTAGAG, CTAGCC, CTAGGA, CTAGGC, GCTAGA, GCTAGC |

Table S4: Over-represented canonical 9-mers per STM topic. Where multiple topics within the same comparison are enriched in the same temperature group, their driving $k$-mers are combined into a single row ($n$ = total count).

| **Domain** | **Comparison** | **Over-rep. in** | $n$ | **Over-represented 9-mers** |
|---|---|---|---|---|
| Bacteria | thermo-meso | Thermophiles | 30 | AGGAGGCCC, AGGCCCAGG, AGGCCCTGG, AGGGCCAGG, AGGGCCTCC, AGGGCCTCG, CAGGGCCTC, CCCAGGCCC, CCCAGGGCC, CCCGGGCCA, CCCTCGAGG, CCCTGGAGG, CCCTGGCCC, CCCTGGCCG, CCTCGAGGC, CCTGGCCCG, CTCGAGGCC, GAGGAGGCC, GAGGAGGGC, GCCAGGGCC, GCCCAGGCC, GCCCAGGGC, GCCCGGGCC, GCCCTGGCC, GCCGAGGCC, GCCTCCTCC, GCCTCGGCC, GCCTGGGCC, GGCCCTGGA, GGGCCTCGA |
| Bacteria | psychro-meso | Psychrophiles | 21 | AAAAAAATA, AAAAAATTA, AAAAATTAA, AAAATAAAT, AAAATATTT, AAAATTAAA, AAAATTATT, AAATAAAAT, AAATTAAAA, AAATTTAAA, AATTAAAAA, AATTTAAAA, AATTTTAAA, ATTTAAAAA, ATTTTAAAA, ATTTTTAAA, ATTTTTTAA, TATTTAAAA, TTAAAAAAA, TTTAAAAAA, TTTTAAAAA |
| Bacteria | thermo-psychro | Thermophiles | 75 | CGCGCCGGC, AAAAAATAT, AAAAGAAAT, AAATTTCTT, AACGAAACG, AACGAATCG, ACGAACGAA, ACGACGGCG, ATCGAACGA, ATCGTTTCG, ATTCGTTCG, ATTTCAAAA, CAAATCGCG, CATCGCTTC, CGAAACGAA, CGACAAACG, CGAGCGCGA, CGATCGAAA, CGATTGAAC, CGCGAGCGC, CGGCGACGA, CGGCGAGCA, CGTCGAGCA, CGTCGCCGC, CGTCGTCGA, GAAACGATC, GAACGATGA, GCGCGCCGC, GCGCGGCGA, GCTCGACGA, GGCGGCGAC, AGGAGGCCC, AGGAGGGCC, AGGCCCTGG, AGGGCCAGG, AGGGCCTCC, CAGGGCCTC, CCCAGGGCC, CCCCCAGGG, CCCTCGAGG, CCCTGGCCC, CCTCGAGGC, CTCGAGGCC, GAGGAGGCC, GGCCCTGGA, AAAAAAGAA, AAAAACAAA, AAAACCGGC, AACCAGGCT, AATACCCGG, AATGGCCAG, ACCGGCAAA, ACTGGCCAG, ATCTGGCCA, ATTGCCCGC, ATTGCCCGG, ATTGCCCTG, ATTGCCGCC, ATTGGCCAG, ATTTCCCGG, CATTGCCGA, CCATTGCCG, CCGGCAAAA, CCGGGATAA, CGCGACTAA, CTGATTGCC, GCAATGGCC, GCATTGGCC, GCCAGGTTA, GCCATTGCC, GCCGGCAAA, GGCAATGCC, GGCGGCAAA, TGCCGCCAA, TGGCGGCAA |

*(continued)*

| Domain | Comparison | Over-rep. in | $n$ | Over-represented 9-mers |
|---|---|---|---|---|
| | | Psychrophiles | 41 | AAAAAATAA, TTTATAAAA, AAAAAAGAA, AAAAAATTA, AAAAAGAAA, AAAAATAAT, AAAAATTTT, AAAAGAAAA, AAAATAATT, AAAGAAAAA, AAGAAAAGA, AAGAAGAGA, AAGAGAGGA, AAGAGATAG, AATAAAAAA, AGAAGAGAG, AGAAGAGAT, AGAGAAGAG, ATCTTCTCA, ATTAAAAAA, CACCTCCTC, CTTCAAAGA, CTTTGAAGA, GAAGAGAGA, GAAGAGATA, GAGAAGAGA, GAGAAGATA, GAGGAGATA, GATAGAAGA, TAAAAGAAA, TATGAGGGA, TCTTCTCAA, AAAATTAAA, AAATTATAA, AAATTTTAA, AATTTAAAA, AATTTTAAA, ATTTTAAAA, TCTTTTAAA, TTTAAAAAA, TTTTAAAAA |
| Archaea | thermo-meso | Thermophiles | 4 | AAGGAGCTC, AGAAGCTCA, AGCTCAAGG, CTCAAGCTC |

### G Sequence Logos for Driving 9-mers

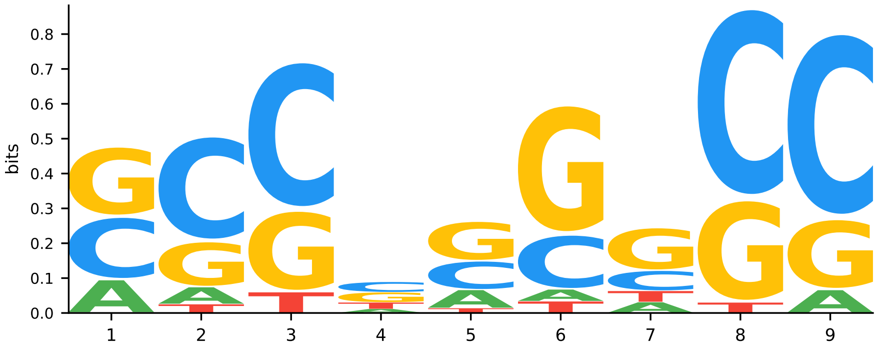


Figure S7: Sequence logo of driving 9-mers overrepresented in bacterial thermophiles (thermo-meso).

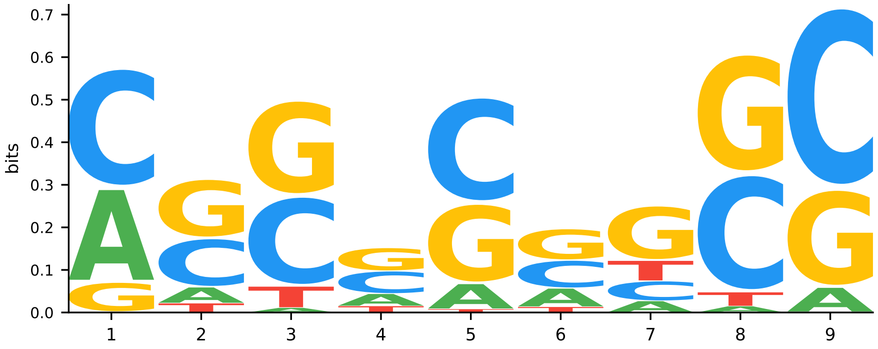


Figure S8: Sequence logo of driving 9-mers overrepresented in bacterial thermophiles (thermo-psychro).

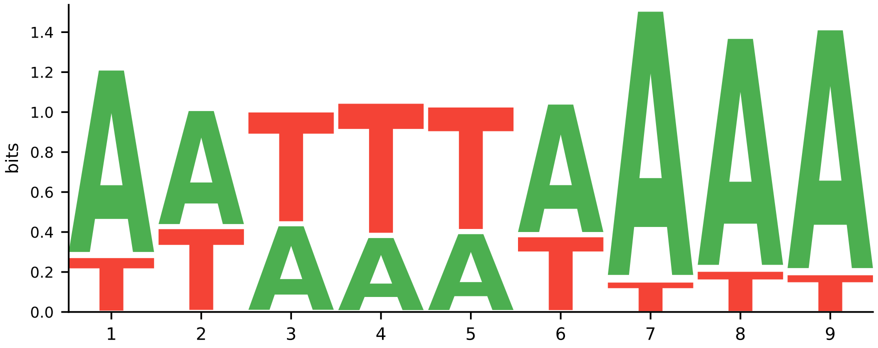


Figure S9: Sequence logo of driving 9-mers overrepresented in bacterial psychrophiles (psychro-meso).

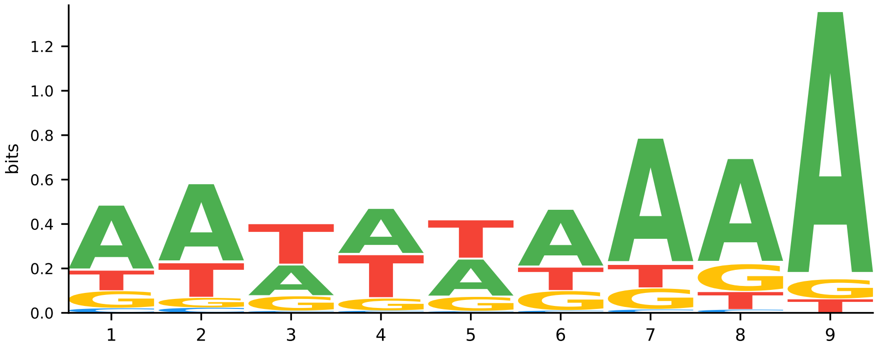


Figure S10: Sequence logo of driving 9-mers overrepresented in bacterial psychrophiles (thermo-psychro).

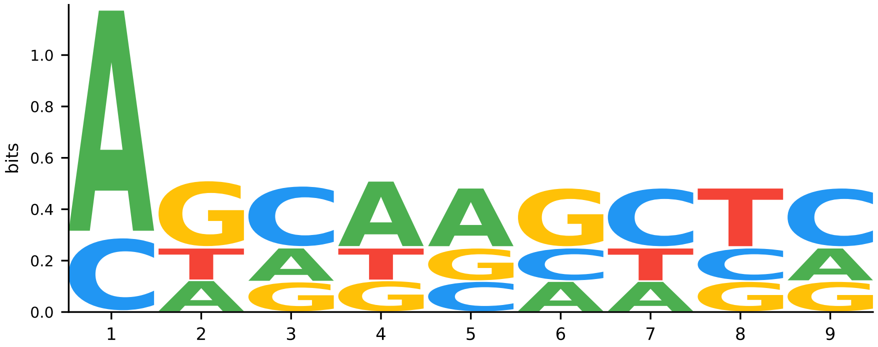


Figure S11: Sequence logo of driving 9-mers overrepresented in archaeal thermophiles (thermo-meso).
